# Fitness-valley crossing with generalized parent-offspring transmission

**DOI:** 10.1101/025502

**Authors:** M.M. Osmond, S.P. Otto

**Affiliations:** Department of Zoology, University of British Columbia, Vancouver, British Columbia V6T 1Z4, Canada

**Keywords:** cultural evolution, cytonuclear inheritance, meiotic drive, peak shift, population genetics, valley crossing

## Abstract

Simple and ubiquitous gene interactions create rugged fitness landscapes composed of coadapted gene complexes separated by “valleys” of low fitness. Crossing such fitness valleys allows a population to escape suboptimal local fitness peaks to become better adapted. This is the premise of Sewall Wright’s shifting balance process. Here we generalize the theory of fitness-valley crossing in the two-locus, bi-allelic case by allowing bias in parent-offspring transmission. This generalization extends the existing mathematical framework to genetic systems with segregation distortion and uniparental inheritance. Our results are also flexible enough to provide insight into shifts between alternate stable states in cultural systems with “transmission valleys”. Using a semi-deterministic analysis and a stochastic diffusion approximation, we focus on the limiting step in valley crossing: the first appearance of the genotype on the new fitness peak whose lineage will eventually fix. We then apply our results to specific cases of segregation distortion, uniparental inheritance, and cultural transmission. Segregation distortion favouring mutant alleles facilitates crossing most when recombination and mutation are rare, i.e., scenarios where crossing is otherwise unlikely. Interactions with more mutable genes (e.g., uniparental inherited cytoplasmic elements) substantially reduce crossing times. Despite component traits being passed on poorly in the previous cultural background, small advantages in the transmission of a new combination of cultural traits can greatly facilitate a cultural transition. While peak shifts are unlikely under many of the common assumptions of population genetic theory, relaxing some of these assumptions can promote fitness-valley crossing.

## 1. Introduction

Epistasis and underdominance create rugged fitness landscapes on which adaptation may require a population to acquire multiple, individually-deleterious mutations that are collectively advantageous. Using the adaptive landscape metaphor, we say the population faces a fitness “valley” (Wright, 1932). Such valleys appear to be common in nature (Weinreich et al., 2005; Szendro et al., 2013, but see Carneiro and Hartl, 2010) and affect, among other things, speciation by reproductive isolation, the evolution of sex, the evolvability of populations, and the predictability of evolution (Szendro et al., 2013). Here we are interested in the speed and likelihood of fitness-valley crossing, which we determine by examining the first appearance of an individual with the collectively advantageous set of mutations whose lineage will eventually spread to fixation.

Believing epistasis to be ubiquitous, Sewall Wright (1931; 1932) formulated his “shifting balance theory”, which describes evolution as a series of fitness-valley crossings. In phase one of the shifting balance process, small, partially-isolated subpopulations (demes) descend into fitness valleys by genetic drift. The new mutations are selected against when rare, as they will tend to occur alone as single deleterious alleles. Eventually drift may allow the deleterious mutations to reach appreciable frequencies in at least one deme. Once multiple synergistically-acting mutations arise together, they begin to be locally favoured by selection. In phase two, these favoured combinations of mutations sweep to fixation, and those demes ascend the new “fitness peak”. Finally, in phase three, the demes that reach the new fitness peak send out migrants whose genes invade and fix in the remaining demes, eventually “pulling” the entire population up to the new fitness peak. Our focus here is in the first appearance of a genotype on the new fitness peak whose lineage will eventually fix, considering a single isolated deme. This is typically the longest stage of phases one and two (Stephan, 1996) and hence is likely the limiting step in fitness-valley crossing.

Fitness-valley crossing has been investigated in a large number of theoretical studies. In the context of multiple loci with reciprocal sign epistasis, the first appearance of the genotype with the best combination of alleles has been the focus of a few studies (Phillips, 1996; Christiansen et al., 1998; Hadany, 2003; Hadany et al., 2004; Weissman et al., 2009, 2010). Many authors have gone on to examine the remainder of phases one and two (Crow and Kimura, 1965; Eshel and Feldman, 1970; Karlin and McGregor, 1971; Kimura, 1985; Barton and Rouhani, 1987; Kimura, 1990; Phillips, 1996; Michalakis and Slatkin, 1996; Stephan, 1996; Weinreich and Chao, 2005; Weissman et al., 2009, 2010), as well as phase three (Kimura, 1990; Crow et al., 1990; Barton, 1992; Kondrashov, 1992; Phillips, 1993; Gavrilets, 1996; Hadany, 2003; Hadany et al., 2004). Similar attention has been given to situations with a single underdominant locus (Slatkin, 1981; Gillespie, 1984; Barton and Rouhani, 1993; Peck et al., 1998) or a quantitative trait (Lande, 1985a; Barton and Rouhani, 1987; Rouhani and Barton, 1987a, b; Charlesworth and Rouhani, 1988; Barton and Rouhani, 1993). The theoretical and empirical support for Wright’s shifting balance process has been summarized and debated (Coyne et al., 1997; Wade and Goodnight, 1998; Coyne et al., 2000; Whitlock and Phillips, 2000; Coyne et al., 2000; Goodnight and Wade, 2000; Goodnight, 2013), and the general consensus appears to be that, unless the valley is shallow (weakly deleterious intermediates), crossing a fitness valley is unlikely.

Despite the abundance of literature on fitness-valley crossing, the above studies all assume perfect Mendelian inheritance. The question therefore remains: how robust are our ideas of fitness-valley crossing to deviations from Mendelian inheritance? Specifically, how does transmission bias (e.g., meiotic drive or uniparental inheritance) affect the speed and likelihood of valley crossing? Departing from strict Mendelian inheritance also allows us to consider the idea of valley crossing in cultures, considering the spread of memes (Dawkins, 1976) rather than genes. This simultaneously adds a level of complexity to current mathematical models of cultural transmission, which typically consider only one cultural trait at a time (e.g., Cavalli-Sforza and Feldman, 1981; but see, e.g., Ihara and Feldman, 2004; Creanza et al., 2012).

Transmission bias in the form of segregation distortion is likely to have a large effect on valley crossing, as distortion represents a second level of selection (Sandler and Novitski, 1957; Hartl, 1970). Insight into how segregation distortion affects valley crossing comes from models of underdominant chromosomal rearrangements (mathematically equivalent to models with *one* diploid biallelic locus), which often find meiotic drive to be a mechanism allowing fixation of a new mutant homokaryotype (Bengtsson and Bodmer, 1976; Hedrick, 1981; Walsh, 1982). Populations that have fixed alternate homokaryotypes produce heterokaryotype hybrids, which have low viability and/or fertility; thus gene flow between these populations is reduced. Segregation distortion is therefore thought to be a mechanism that promotes rapid speciation (stasipatric speciation; White, 1978). Although the role of underdominance in chromosomal speciation has recently been questioned (reveiwed in Rieseberg, 2001; Hoffmann and Rieseberg, 2008; Faria and Navarro, 2010; Kirkpatrick, 2010), it is hypothesized to be relevant in annual plants (Hoffmann and Rieseberg, 2008) and appears to play a dominant role in maintaining reproductive isolation in sunflowers (Lai et al., 2005) and monkey flowers (Stathos and Fishman, 2014).

Another common form of transmission bias is sex specific, with the extreme case being uniparental inheritance. In genetic transmission, strict uniparental inheritance is common for organelle genomes, such as the mitochondria, which is typically inherited from the mother. Uniparental inheritance will tend to imply further asymmetries. For instance, the mutation rate of mitochondrial genes is estimated to be two orders of magnitude larger than the mutation rate of nuclear genes in many animals (e.g., Linnane et al., 1989). Higher mutation rates will likely facilitate crossing. That said, higher mutation rates in only one gene may have limited effect because the production of double mutants by recombination will be constrained by the availability of the rarer single mutant. Previous models of fitness-valley crossing have tended to ignore asymmetries (but see Appendix C of Weissman et al., 2010).

Transmission bias is an integral characteristic of cultural transmission, where it is referred to as “cultural selection” (Cavalli-Sforza and Feldman, 1981; Boyd and Richerson, 1985). However, to the best of our knowledge, no attempts have been made to examine the evolution of cultural traits (memes) in the presence of a “fitness” valley. Boyd (2001) reviews the genetic theory of the shifting balance, and notes that it could be applied to culture, but no explicit cultural models were presented. Meanwhile, instances such as the so-called “demographic transition” in 19^*th*^ century western Europe, where societies transitioned from less educated, large families to more educated, small families (Borgerhoff Mulder, 1998), suggest that alternate combinations of cultural traits (e.g.,‘value of education’ and ‘family-size preference’) can be stable and that peak shifts may occur in cultural evolution. In fact, alternate stable cultural states may be pervasive (Boyd and Richerson, 2010), as alluded to by the common saying that people are “stuck in their ways.” Paradigm shifts in the history of science (Kuhn, 1962) may provide further examples (Fog, 1999). Cultural peak shifts can also be relatively trivial; for instance, changing the unit of time from seconds, minutes, and hours to a decimal system is only advantageous if we also change units that are based on seconds, such as the joule and volt (Fog, 1999).

Here we focus on a population genetic model with two bi-allelic loci under haploid selection in a randomly-mating, finite population. This model can easily be reduced to a single-locus model with two alleles and diploid selection, which is formally equivalent to a model of chromosomal rearrangements (e.g., a chromosome has an inversion or not). Interpreting genes as memes produces a model of vertically-transmitted cultural evolution. Our model incorporates both transmission bias and asymmetries in mutation and initial numbers of single mutants. We first give a rough semi-deterministic sketch to develop some intuition, then follow with a stochastic analysis using a diffusion approximation. Our analysis corresponds to the stochastic simultaneous fixation regime of Weinreich and Chao (2005), and the neutral stochastic tunnelling and deleterious tunnelling regimes of Weissman et al. (2010), where the appearance of the new, favourable, and eventually successful “double mutant” occurs before the fixation of the neutral or deleterious “single mutants”. Finally, we apply our results to the specific cases of segregation distortion, uniparental inheritance, and cultural transmission.

We derive the expected time until the appearance of a double mutant whose lineage will fix when single mutants are continuously generated by mutation from residents (the stochastic model assumes neutral single mutants). We also use the stochastic model to derive the probability that a double mutant appears and fixes given an initial stock of deleterious single mutants that is not replenished by mutation. Given typical per-locus mutation rates, valley crossing is generally found to be a slow and unlikely outcome under fair Mendelian transmission, even when single mutants are selectively neutral. Segregation distortion, in favour of wild-type or mutant alleles, affects crossing most when recombination and mutation are rare, the scenarios where crossing is otherwise unlikely. Cytonuclear inheritance allows increased mutational asymmetries between the two loci; higher mutation rates lead to more single mutants and hence faster valley crossing, but, when holding the average mutation rate constant, asymmetries hinder crossing by reducing the probability that the single mutants recombine to produce double mutants. Finally, we show that, when new cultural ideas or practices are not too poorly transmitted when arising individually within the previous cultural background, a transmission advantage of the new combination greatly facilitates cultural transitions.

## 2. Model and Results

Consider two loci, **A** and **B**, with *x*_*ij*_ the current frequency of genotype *A*_*i*_*B*_*j*_, where *i*, *j* ∊{1, 2, *…,p*} are the alleles carried by the individual. When an *A*_*i*_*B*_*j*_ individual mates with an *A*_*i*_*B*_*l*_ individual, they produce an *A*_*m*_*B*_*n*_ offspring with probability 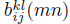. Summing over all possible offspring types 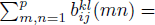 We can specify that the bottom index (here *kl*) denotes the genotype of the mother, while the top index (here *kl*) denotes the genotype of the father. As a consequence, transmission biases according to parental sex 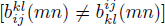 are allowed. When considering sex-biased transmission we assume the frequencies *x*_*ij*_ are the same in females and males (i.e., no sex linkage and no sex-based differences in selection), which is automatically the case in hermaphrodites.

Random mating and offspring production is followed by haploid viability selection, which occurs immediately before censusing. The population size, *N*, is constant and discrete, and generations are non-overlapping. Then the expected frequency of *A*_*m*_*B*_*n*_ in the next generation, 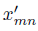, solves

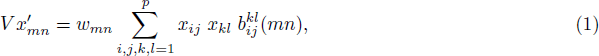

where *w*_*mn*_ ≥ 0 is the relative viability of *A*_*m*_*B*_*n*_ and *V* is the sum of the right hand side of Equation (1) over all genotypes, which keeps the frequencies summed to one.

Denote the probability that a mating between an *A*_*i*_*B*_*j*_ mother and an *A*_*k*_*B*_*l*_ father produces an *A*_*m*_*B*_*n*_ offspring that survives one round of viability selection by 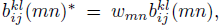 where the asterisk indicates “after selection”. And let the average probability that a mating produces *A*_*m*_*B*_*n*_, regardless of which parent was which, be 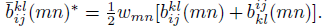 Then (as we will see below) selection on *A*_*i*_*B*_*j*_ in a population of “residents” (*A*_1_*B*_1_) is described by 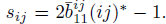 Letting *w*_*ij*_ = 1 + *d*_*ij*_ > 0 describe viability selection and 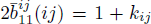 describe transmission bias (− 1 ≤ *k*_*ij*_ ≤ 1), then *s*_*ij*_ = (1 + *d*_*ij*_)(1 + *k*_*ij*_) − 1. Here we define the relative fitness of genotype *A*_*i*_*B*_*j*_ as 1 + *s*_*ij*_, which is determined by both viability and transmission. Thus defined, fitness is a measure of the “transmissibility” of a genotype as it includes several processes (e.g., viability, meiotic drive, recombination, mutation) that affect the number of offspring of a given genotype produced by a parent of that genotype. We will see that it is transmissibility that determines the dynamics of valley crossing.

Without mutation or recombination, fair transmission implies 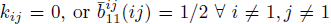. In words, with fair transmission we expect half of all offspring from matings between A_1_B_1_ and *A*_*i*_*B*_*j*_ to be of parental type *A*_*i*_*B*_*j*_. Sex-based inheritance is expected to arise in the form of 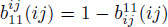 [e.g., maternal inheritance implies 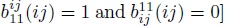, which does not directly impose selection as *k*_*ij*_ = 0. Segregation distortion can, however, impose selection. For example, ignoring mutations, if the *A*_2_ allele is more likely to be transmitted than the *A*_1_ allele (in a *B*_1_ background) we would have 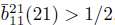, giving *k*_21_ > 0. Interpreting genes as memes, transmission bias *k*_*ij*_ determines the strength of “cultural selection” *(sensu* Cavalli-Sforza and Feldman, 1981) on meme combination *A*_*i*_*B*_*j*_. Previous work on multi-locus peak shifts has assumed that bias does not influence selection (*k*_*ij*_ = 0) and that maternal and paternal types are equally transmitted 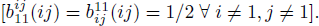

Here we focus on bi-allelic loci (*p* = 2). We are specifically interested in the case where, in a population composed entirely of residents, “single mutants” (*A*_2_*B*_1_ and *A*_1_*B*_2_) are selected against while “double mutants” (*A*_2_*B*_2_) are selectively favoured: *s*_21_, *s*_12_ < 0 < *s*_22_.

Given that the population is composed primarily of residents, with no double mutants as of yet, the population faces a fitness valley. The valley can be created by differences in viability alone, or it can be created by differences in transmission, or both. Here we focus on the limiting step in the peak-shift process, the probability and expected time until a double mutant arises whose lineage will eventually fix. Following the lead of Christiansen et al. (1998), we begin by developing a rough semi-deterministic analysis to gain intuition. A stochastic analysis follows. Table 1 provides a summary of notation and a supplementary *Mathematica* file gives a more detailed derivation of the results.

**Table 1:** Parameters used throughout text

| Symbol | Description |
| --- | --- |
| $x_{ij}$ | frequency of $A_i B_j$ in the current generation |
| $x'_{ij}$ | expected frequency of $A_i B_j$ in the next generation |
| $w_{ij}$ | viability of $A_i B_j$ relative to viability of $A_1 B_1$ |
| $V$ | normalizing factor |
| $N$ | number of individuals in the population |
| $t$ | time, in units of generations |
| $b_{ij}^{kl}(mn)$ | probability $A_i B_j$ mother and $A_k B_l$ father produce $A_m B_n$ offspring |
| $\bar{b}_{ij}^{kl}(mn)^*$ | average probability of surviving $A_m B_n$ offspring from $A_i B_j \times A_k B_l$ mating,<br>$\frac{1}{2}w_{mn}[b_{ij}^{kl}(mn) + b_{kl}^{ij}(mn)]$ |
| $\mu_{ij}^{kl}(mn)^*$ | probability of surviving *mutant* offspring, $\bar{b}_{ij}^{kl}(mn)^*$, $m \notin \{i, k\}$, $n \notin \{j, l\}$ |
| $r_{ij}^{kl}(mn)^*$ | probability of surviving *recombinant* offspring, $\bar{b}_{ij}^{kl}(mn)^*$, $m \in \{i, k\}$, $n \in \{j, l\}$,<br>$mn \notin \{ij, kl\}$ |
| $\bar{b}_{ij}^{kl}(mn)$ | average transmission probability before selection, $\bar{b}_{ij}^{kl}(mn)^*/w_{mn}$<br>[similarly for $\mu_{ij}^{kl}(mn)$ and $r_{ij}^{kl}(mn)$] |
| $s_{ij}$ | selection on $A_i B_j$ in a resident population, $2\bar{b}_{11}^{ij}(ij)^* - 1$ |
| $T$ | generations until first successful double mutant arises |
| $u_{22}$ | probability that a double mutant begins a lineage that will fix |
| $i_t$ | number of $A_2 B_1$ individuals in generation $t$ (similarly for $A_1 B_2$, $j_t$) |
| $X(t)$ | numbers of single mutants in generation $t$ assuming no double mutants, $(i_t, j_t)$ |
| $\Delta i$ | change in number of $A_2 B_1$ individuals, $i_{t+1} - i_t$ (similarly for $A_1 B_2$, $\Delta j = j_{t+1} - j_t$) |
| $\alpha, \beta$ | scaling parameters in diffusion process |
| $\tau$ | scaled unit of time, $t/N^\alpha$ |
| $Y(\tau)$ | scaled frequency of $A_2 B_1$, $i_\tau/N^\beta$ (similarly for $A_1 B_2$, $Z(\tau) = j_\tau/N^\beta$) |
| $\mu_Y(y)$ | first moment of $\Delta Y = Y(\tau + 1) - Y(\tau)$ given $Y(\tau) = y = i/N^\beta$ (similarly for $Z$) |
| $\sigma_Y^2(y)$ | second moment of $\Delta Y$ given $Y(\tau) = y$ (similarly for $Z$) |
| $\kappa(y, z)$ | rate diffusion killed by successful double mutants given $Y(0) = y, Z(0) = z$ |
| $B_{ij}^{kl}(mn)$ | scaled transmission probability, $b_{ij}^{kl}(mn)N^\beta$ |
| $R_{21}^{12}(22)$ | scaled (rare) recombination from single mutants to double mutants, $r_{21}^{12}(22)N^{1/2}$ |
| $S_{ij}$ | scaled selection on $A_i B_j$ in population of residents, $s_{ij}N^\beta$ |
| $\tilde{T}(y, z)$ | scaled time until first successful double mutant given $Y(0) = y, Z(0) = z, TN^\alpha$ |
| $m$ | index for single mutant types when equivalent (e.g., $s_m = s_{21} = s_{12}$) |
| $c_\mu$ | mutation rate at locus **B** relative to locus **A**, $\nu/\mu$ |
| $c_n$ | initial number of $A_1 B_2$ individuals, relative to $A_2 B_1$, $j_0/i_0$ |
| $\xi$ | scaled frequency of single mutants ($y + z$ or $c_i y = z$, depending on assumptions) |
| $u(y, z)$ | probability no successful double mutant appears given $Y(0) = y, Z(0) = z$ |
| $n_0$ | initial number of single mutants, $i_0 + j_0$ |

### 2.1. *Semi-deterministic analysis*

#### 2.1.1 *Single mutant dynamics*

Selection against single mutants keeps their frequencies (*x*_21_ and *x*_12_) small. Let these frequencies be proportional to some small number ε ≪ 1. Let the probability that an offspring inherits an allele that neither parent possesses [i.e., mutation; e.g., 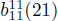] be of the same small order ∊. Then, for large *N*, the frequencies of the single mutants in the next generation are

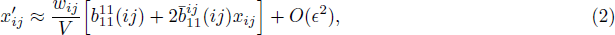

where *i* ≠ *j* and *O*(∊^2^) captures terms of order ∊^2^ and smaller.

We will write 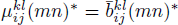 when *m* ∉ {*i*, *k*} or *n* ∉ {*j*, *l*} to highlight the fact that a mutation has occurred. Then, ignoring O(∊^2^), the frequencies of single mutants, which are initially absent [*x*_*ij*_(0) = 0], in generation *t* are

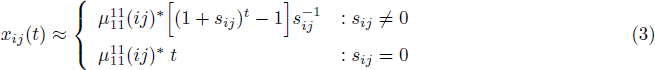

Viability and transmission are thus coupled together (in *s*_*ij*_) throughout our results, and it is primarily the total amount of selection on *A*_*i*_*B*_*j*_ in a population of residents (*s*_*ij*_) that determines the dynamics. [As a technical aside, this is not true in the first generation that mutants appear, via 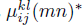, but this is simply because of the order of the life cycle chosen, where these mutants experience viability selection, but not transmission biases, when they first occur.]

Equation (3) assumes the normalizing factor *V* remains near 1 over the *t* generations, which is the case when single mutants are rare, as is generally true when single mutants are selected against, *s*_*ij*_ < 0 ∀ *i* ≠ *j*. When *s*_*ij*_ < 0 and there has been a sufficiently long period of selection, t > − 1/*s*_*ij*_, the single mutant frequencies approach mutation-selection balance 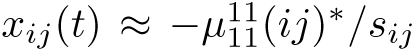 This assumes the probability of mutation, 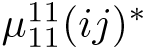, is small relative to the strength of selection, *s*_*ij*_. We next derive a semi-deterministic solution for the crossing time, *T*, given mutation-selection balance is reached. In Appendix A we follow Christiansen et al. (1998) to derive the semi-deterministic crossing time when crossing occurs before mutation-selection balance is reached; this occurs when −*s*_*ij*_T ≪ 1, which can only be the case if the valley is shallow, − *s*_*ij*_ ≪ 1.

#### 2.1.2 *Waiting time for first successful double mutant*

We now turn to calculating the waiting time until a double mutant that is able to establish first arises. Assume the probability two residents mate to produce a double mutant (i.e., a double mutation), 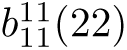, is very rare, on the order of ∊^2^. Then the expected frequency of double mutants in the next generation before selection, assuming single mutant are rare and there are currently no double mutants *x*_22_ = 0, is

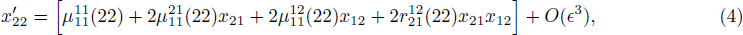

where we write 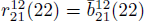 to highlight the fact that a double mutant has effectively been produced by recombination. The expected frequency of double mutants (Equation 4) is measured before viability selection to avoid artificially adjusting the double mutant frequency by its viability difference before it appears.

In a truly deterministic model (N → ∞) double mutants are present at frequency 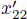 after a single bout of reproduction. However, assuming no double mutants have yet appeared, we can use *x*_22_(*t*) as a rough approximation for the probability of a double mutant first arising in generation _t_ (Christiansen et al., 1998). Summing *t* from 0 to *t*′ gives the cumulative probability of observing a double mutant in any of the *t*′ generations. The generation *T*′ at which the cumulative probability reaches 1/*N* can be used as an estimate of the time we expect to wait until the first double mutant has arisen (Christiansen et al., 1998).

Here we are more interested in the waiting time until the first *successful* double mutant appears (i.e., one whose lineage will eventually fix). We therefore want to multiply the probability that a double mutant appears at time *t*, *x*_22_(*t*), by the probability it will fix before taking the sum over t. Using Kimura’s (1962) approximation, the probability a double mutant fixes is

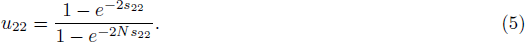

With a weak double mutant advantage, 0 < *s*_22_ ≪ 1, in a large population, *Ns*_22_ ≫ 1, Equation (5) simplifies to the familiar 2*s*_22_ (Haldane, 1927).

The selection coefficient *s*_22_ can be calculated from the number of double mutant offspring a newly arisen double mutant is expected to leave in the next generation, given that the mean number of offspring per individual is one, such that the population size is constant. This expectation, 1 + *s*_22_, is the probability of mating with a given type, multiplied by the probability of producing a double mutant offspring, multiplied by the probability of surviving to the next generation, summed over all possible matings

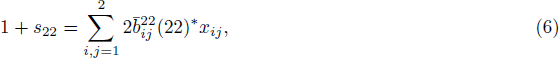

where *x*_22_ = 0 in the remaining population (i.e., the double mutant does not mate with itself). Without transmission bias, mutation, or recombination, 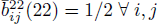 and Equation (6) reduces to the familiar *s*_22_ = *w*_22_ − 1. Here we allow bias, mutation, and recombination, and assume single mutants are sufficiently rare, giving 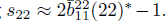 This implies that selection on the double mutant (including transmission) is constant over time and that fixation depends only on its dynamics in a population composed almost entirely of residents. With recombination and otherwise fair transmission we have 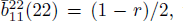 where *r* is the probability of recombination between a double mutant and a resident. Writing *w*_22_ = 1 + *s* and assuming both *s* and *r* are small, recovers the well-known first-order approximation *s*_22_ = *s* − *r* (Crow and Kimura, 1965). This expression highlights the fact that recombination can reduce the probability of fixation by breaking up favourable gene combinations (Crow and Kimura, 1965).

When selection is strong and mutation is rare, relative to the strength of genetic drift, the time to fixation is dominated by the time to the arrival of a successful mutant (Gillespie, 1984; Weinreich and Chao, 2005; Weissman et al., 2010). The waiting time until the first successful double mutant, which we derive below, therefore well approximates the fixation time of a double mutant within a population when double mutants are advantageous but rarely produced, 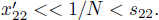

*Crossing time given mutation-selection balance.* When enough time has passed (*t* > −1/*s*_*ij*_) the singlemutant frequencies approach mutation-selection balance (MSB), 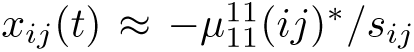 Using these frequencies in Equation (4) gives the expected frequency of double mutants in the next generation, which does not change until a double mutant arises, i.e., 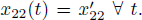 Summing 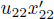 over T_MSB_ generations, setting equal to 1/*N*, and solving for T_MSB_ gives an estimate of the number of generations we expect to wait for a successful double mutant to arise when beginning from mutation-selection balance

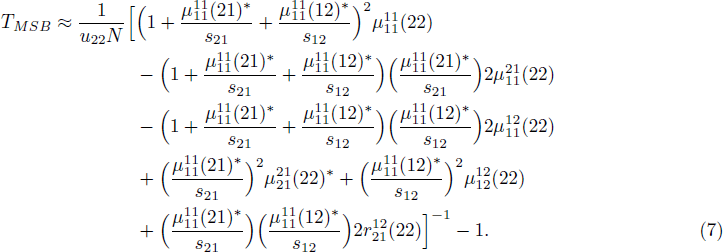

In our numerical examples, we will track the waiting time until a successful double mutant arises in a population that has recently established and is fixed for the resident type (e.g., following a bottleneck or a founder event). This time can be approximated by the time that it takes to reach mutation-selection balance, *T*_0_, and the establishment time once there

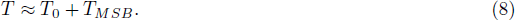

Here we use 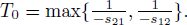 As the deleterious single mutants approach neutrality (*s*_*ij*_ → 0^−^∀ *i* ≠ *j*) the waiting time *from* mutation-selection balance, T_MSB_, decreases (because there are more single mutants segregating), but the waiting time *to* mutation-selection balance, T0, increases dramatically because it takes longer to produce the higher segregating frequencies of single mutants. As − *S*_*j*_ becomes small enough such that T < − 1/*s*_*ij*_ the approximation breaks down and we must use the non-equilibrium solution derived in Appendix A.

With symmetric Mendelian assumptions, weak selection on single mutants (δ=1 − *w*_*ij*_∀*i*≠*j*), rare mutation (*μ*), and infrequent recombination [such that *u*_*f*_ ≈ 2(*s* − *r*) ≈ 2*s*], the rate of production of successful double mutants from mutation selection balance (Equation 7) is

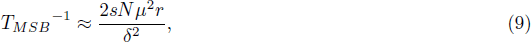

aligning with equation 4 in Weissman et al. (2010, see supplementary *Mathematica* file). This result preforms well when T_MSB_-^1^ < (**δ**, or, equivalently, when 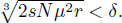

### 2.2. *Stochastic analysis*

#### 2.2.1 *Markov process*

Fitness-valley crossing is naturally a stochastic process. We thus now consider the Wright-Fisher model, where the next generation is formed by choosing *N* offspring, with replacement, from a multinomial distribution with frequency parameters 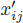(Equation 1). Let the number of *A*_2_*B*_1_ and *A*_1_*B*_2_ single mutants in generation t be it and jt, respectively. Given that there are currently no double mutants, we have *N* − *i*_*t*_ − *j*_t_ resident individuals and we let *X*(*t*) = (*i*_*t*_, *j*_*t*_) describe the state of the system in generation *t*. Let the expected frequencies in the *t* + 1 generation, conditional on *X*(*t*) = (*i*,*j*), be 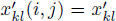, with *x*_22_ = 0 The transition probabilities to states without double mutants are the

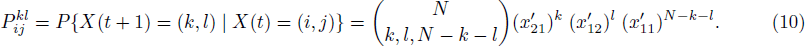

Note that summing over all *k*, *l* ∈ {0,1,…, *N*} gives 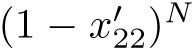, the probability that no double mutant is sampled. Equation (10) describes a sub-stochastic transition matrix for the Markov process.

Next, let *H* be the state with any positive number of double mutants. We then have the transition probabilities 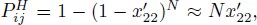 where the approximation assumes a small expected frequency of double mutants in the next generation, 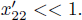. To calculate the waiting time until the first *successful* double mutant, we replace 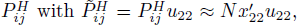 ignoring the segregation of double mutants when lost. *H* is now the state with any positive number of successful double mutants. Dividing each 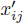 in Equation (10) by the probability a double mutant does not arise 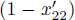 and multiplying by the probability a double mutant does not arise and fix 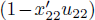 ensures the columns sum to one. To complete the transition matrix we make *H* an absorbing state: 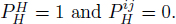

We can describe this process, in part, by the moments for the change in number of single mutants, conditional on the process not being killed by a successful double mutant. The *n*^*th*^ moment for the change in the number of *A*_2_*B*_1_ individuals, 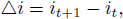 is

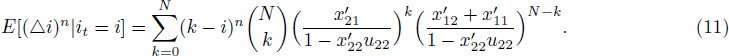

Similar equations can be computed for the change in the number of *A*_1_*B*_2_ individuals, Δ*j* = *j*_*t*+1_ − *j*_*t*_.

To make analytic progress we use the moment equations to approximate the Markov chain with a diffusion process (Karlin and Taylor, 1981, Ch. 15). We do so by taking the large population limit (N → ∞) while finding the appropriate scalings to ensure finite drift and diffusion terms (Appendix B).

### 2.2.2 *Crossing time with neutral single mutants*

The diffusion process yields a partial differential equation describing the expected time until a successful double mutant arises given that we begin with *N*^*β*^*y* individuals of type *A*_2_*B*_1_ and *N*^*β*^*z* individuals of type *A*_1_*B*_2_ (Christiansen et al., 1998)

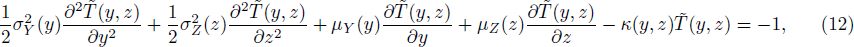

where 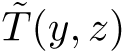 refers to time scaled in units of *N*^β^ generations (parameters defined in Table 1 and Appendix B). In Appendix C we solve Equation (12) under the two scenarios explored in Christiansen et al. (1998): with and without recombination from neutral single mutants to double mutants when the population begins with only residents, but here generalized to allow unequal mutation rates and sex-biased transmission. While the neutrality assumption precludes the existence of a fitness valley, it provides a minimum for the expected time to observe a successful double mutant. Previous studies have suggested that fitness valleys will only be crossed if single mutants are nearly neutral (e.g., Walsh, 1982).

### 2.2.3 *Probability of crossing from standing variation*

The diffusion process can also be used to describe the production of successful double mutants from an initial stock of single mutants (i.e., evolution from standing variation). Specifically, assuming that residents don’t mutate 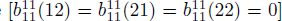 the process has two absorbing states, fixation of *A*_1_*B*_1_ and fixation of *A*_2_*B*_2_ (a successful double mutant appears and the process is killed). The probability of fixation of residents is the solution, *u*(*y*, *z*), of (Karlin and Taylor, 1981)

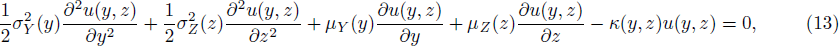

with terms defined in Appendix B. The probability that a successful double mutant arises is therefore 1 − *u*(*y*, *z*). Karlin and Tavare (1981) used a similar equation to find the probability of detecting a lethal homozygote in the one locus, diploid case with Mendelian transmission.

*Deleterious single mutants without recombination.* With no recombination from single mutants to double mutants 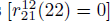 we have scaling parameter *β* = 1/2. Then, with equal selection on single mutants and some mutational symmetry between the two loci (see supplementary *Mathematica* file), the single mutants are equivalent and we can concern ourselves with only their sum ξ = *y* + *z*. Equation (13) then collapses to

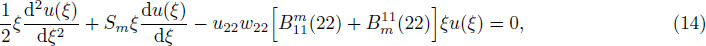

where *S*_*m*_= *s*21*N*^*β*^ = *s*_12_*N*^*β*^ is scaled selection on single mutants and 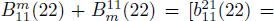 is the scaled mutation probability from single mutants to double mutants.

The boundary conditions are *u*(0) = 1 and *u*(∞) = 0. Solving the boundary value problem gives the probability of a double mutant appearing when starting with *n*_0_ = *i*_0_ + *j*_0_ single mutants

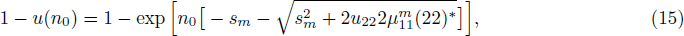

where *s*_*m*_ = *s*_21_ = *s*_12_ is the total strength of selection on each single mutant type. Setting *n*_0_ = 1 gives the probability a newly arisen single mutant will begin a lineage which eventually produces a successful double mutant.

Interestingly, Equation (15) does not depend strongly on population size, *N*. Without recombination double mutants are primarily produced by mutations from single mutants, which are rare and hence always mate with one of the large number of residents. In other words, the production of *A*_2_ and *B*_2_ alleles does not rely on the number of residents but only on the dynamics of the rare single mutants.

*Deleterious single mutants with recombination.* Finally, we examine the probability of a successful double mutant appearing when there is recombination between deleterious single mutants 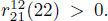 With sufficiently strong selection against single mutants the single mutant frequencies scale as *c*_*n*_*y* ≈ *z* when we begin with initial frequencies *c*_*n*_*y*(0) = *z*(0) and both single mutants are under the same selection pressure, S_21_ = S_12_. Then, without mutation from residents to single mutants, Equation (13) collapses to

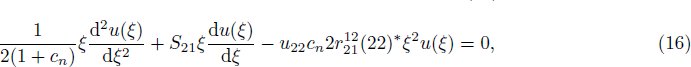

where ξ = *c*_n_ *y* = *z*.

With boundary conditions *u*(0) = 1 and *u*(∞) = 0 the probability of valley crossing is

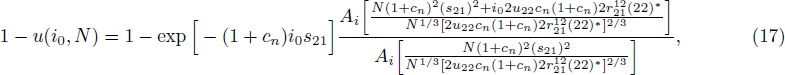

where *A*_*i*_ is the Airy function. Equation (17) extends the one-locus diploid result with Mendelian transmission (equation 28 in Karlin and Tavare, 1981) by allowing unequal single mutant frequencies (*c*_*n*_ ≠ 1) while also incorporating transmission bias, recombination, and double mutant fitness. Equation (17) wellapproximates the Mendelian simulation results of Michalakis and Slatkin (1996, see supplementary *Mathematica* file).

When (*s*_21_)^2^ and *i*_0_ are small, we have the first order approximation

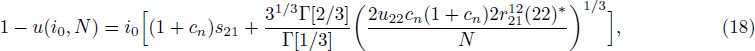

which is valid only when the term in the large square brackets is positive. Equation (18) can be used to show that when holding the initial number of single mutants, (1 + *c*_*n*_)*i*_0_, constant, the probability the double mutant fixes is maximized when there are equal numbers of single mutants, *c*_*n*_ = 1. This is because recombination is most efficient in creating double mutants when single mutants are equally frequent.

### 2.3. *Three scenarios*

We next apply our results to three different scenarios: segregation distortion, cytonuclear inheritance, and cultural transmission.

#### 2.3.1 *Segregation distortion*

One form of segregation distortion, found in the heterothallic fungi *Neurospora intermedia*, is autosomal killing (Burt and Trivers, 2006). In heterozygotes, the presence of a “killer” allele results in the death of a proportion of the spores that contain the wild-type (“susceptible”) allele, leading to a (1 + *k*)/2 frequency of the killing allele at fertilization, 0 < k ≤ 1. Letting *A*_2_ and *B*_2_ represent the killing alleles, and, for the sake of exploration, assuming that cells with one killing allele are functionally equivalent to cells with two, the transmission probabilities are shown in Table 2. The Mendelian case is given by *k* = 0. When −1 ≤ *k* < 0 the allele identities are reversed: *A*_1_ and *B*_1_ are killers and *A*_2_ and *B*_2_ are susceptibles.

**Table 2:**
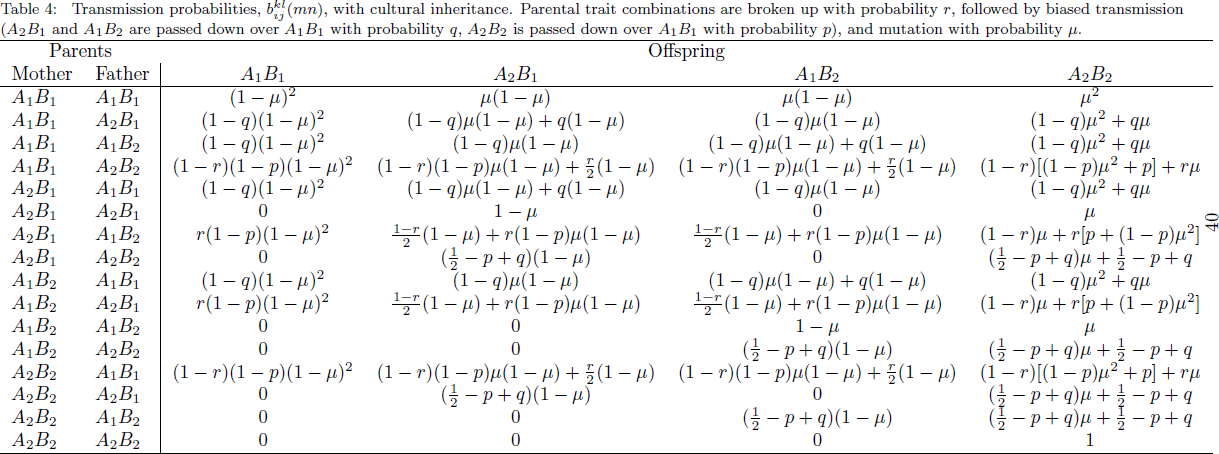
Transmission probabilities, 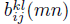, with segregation distortion (autosomal killing). Recombination occurs with probability *r*, followed by autosomal killing of strength 0 ≤ k ≤ 1, and mutation with probability µ. When k > 0 the killing alleles are the mutant alleles (A2 and *B*_2_) and when k < 0 the killing alleles are the resident alleles (*A*_1_ and *B*_1_).

Since segregation distortion imposes selection on single mutants, we can only investigate the effect of segregation distortion on valley crossing with the semi-deterministic crossing time estimates allowing selection on single mutants (Equations 8 and *A*3) and with the crossing probability estimates from standing variation (Equations 15 and 17).

Figure 1 shows the crossing time as a function of the probability of recombination, and how segregation distortion affects this time. Simulations (X’s) well match the numerical solution (Equation A1; dots) and the MSB approximation (Equation 8; solid curves in top panel) over the range of parameters tested. When valley crossing occurs before reaching MSB (bottom panel) a transition occurs between when mutation drives crossing (dashed line; Equation A2) and when recombination does (solid curves; Equation A3), here approximately *r* ≈ 10^−4^. The largest effect of segregation distortion occurs when the crossing time is long, the scenario in which single-mutants must persist the longest before a successful double mutant appears. In addition, observe that as the probability of recombination, *r*, increases above a critical value such that *s*_22_ < 0, the double mutant is broken apart faster than its selective advantage and valley crossing takes longer [equations B21 and B25 in Weissman et al., 2010 approximate the crossing times with no segregation distortion (*k* = 0) when *s*_22_ < 0; see also Lynch, 2010; Altland et al., 2011].

**Figure 1:**
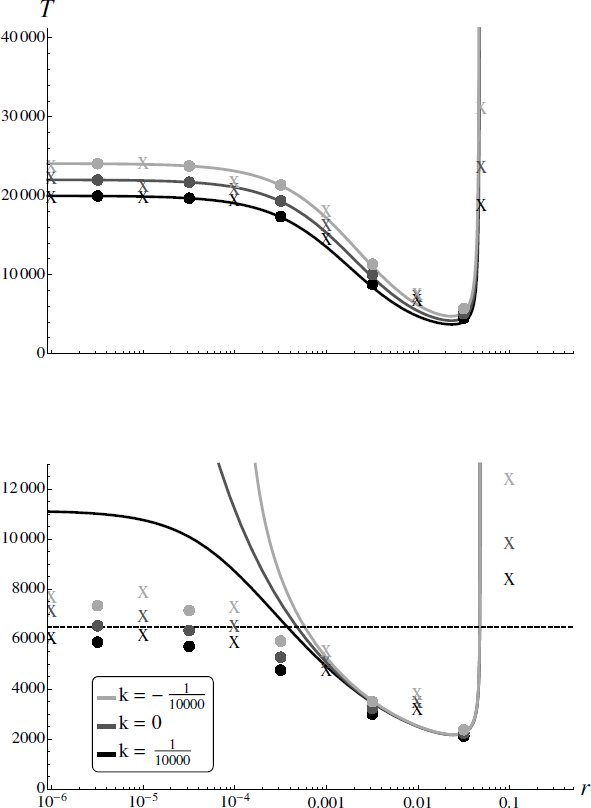
Expected number of generations until a double mutant begins to fix, *T*, as a function of the probability of recombination, *r.* The dots show the full semi-deterministic solution (numerical solution to Equation A1, including higher order terms, allowing both recombination and mutation to generate double mutants). The solid curves show the semi-deterministic results when (top) mutation-selection balance is first reached (Equation 8) and (bottom) mutation-selection balance is not reached and crossing can occur by recombination (Equation A3). The dashed line gives the crossing time when crossing occurs by mutation only, before mutation selection balance is reached, and single mutants are selectively neutral (Equation A2). The X’s are mean simulation results (Appendix D). The grayscale corresponds to *(dark*) distortion favouring single mutants, *k* = 10^−4^; (*medium*) the Mendelian case, *k* = 0; and (*light*) distortion favouring wild-type, *k* = −10^−4^. Parameters: *µ* = 5 x 10^−7^, N = 106, *W22* = 1.05, and (**top**) *w*_21_ = *w*_12_ = 1 − 10^−3^ and (**bottom**) *w*_21_ = *w*_12_ = 1 − 10^−5^.

Figure 2 shows the probability of crossing from standing variation. Again, segregation distortion has a large effect when mutation (top panel) and recombination (bottom panel) are rare, the conditions under which single mutants must persist the longest before a successful double mutant is formed. When crossing occurs by recombination our analytical approximation (Equation 17) overestimates the probability of crossing (bottom panel), especially when the initial number of single mutants is small and therefore subject to strong stochasticity (results not shown). This occurs because the assumption that the ratio of single mutant frequencies in these simulations remains roughly *c*_*n*_ = 1 is violated, reducing the probability that double mutants are formed by recombination.

**Figure 2:**
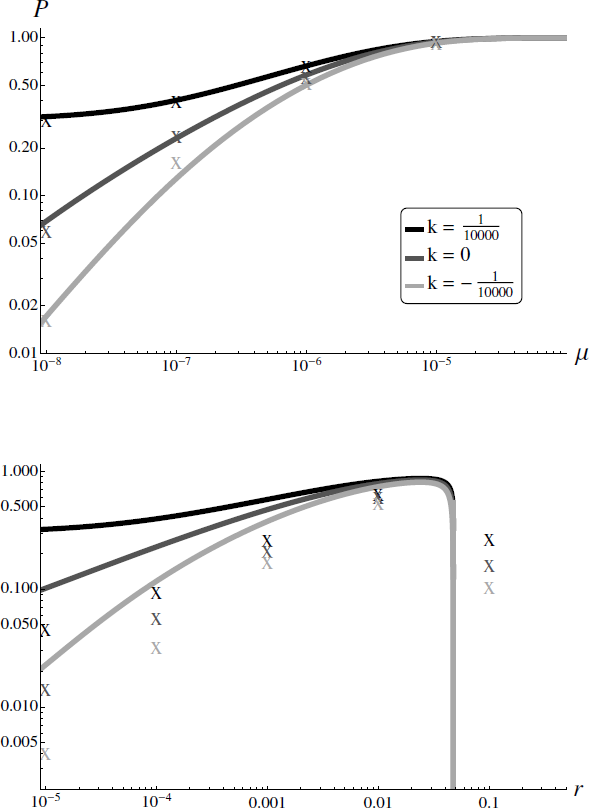
The probability, P = 1 − *u*, of crossing the valley given an initial stock of single mutants (with no further mutations from resident-resident matings) as a function of the rate at which single mutants produce double mutants **(top)** without recombination (Equation 15) and **(bottom)** by recombination only (Equation 17). The X’s are simulation results (Appendix D). Parameters and grayscale as in Figure 1 **(bottom)** with *i*_0_ = *j*_0_ = 1000.

*2.3.2. Cytonuclear inheritance*

We next explore how fitness-valley crossing is affected by uniparental inheritance of one of the traits. This might occur if, for example, there was reciprocal sign epistasis between cytoplasmic and nuclear loci. Without loss of generality we assume that the B trait is always inherited from the mother. For simplicity we assume individuals are hermaphroditic. Here we can use only those results that allow recombination (Equations 8, A3, C5, and 17), as cytoplasmic and nuclear elements are expected to be inherited independently (i.e., 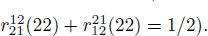

One likely implication of uniparental transmission is asymmetric mutation probabilities. For instance, in animals the mitochondrial mutation rate is two orders of magnitude larger than typical nuclear rates (Linnane et al., 1989). Let µ be the mutation probability in the biparentally inherited A trait and ν be the mutation probability in the uniparentally inherited B trait, with c_*µ*_ = ν/µ the ratio of uniparental to biparental mutation probabilities. The transmission probabilities are shown in Table 3.

**Table 3:**
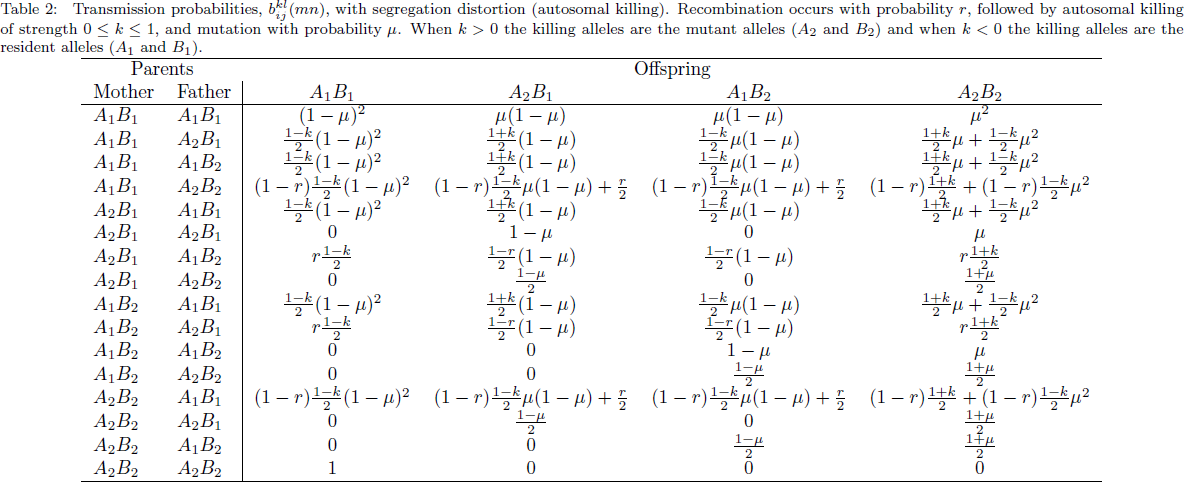
Transmission probabilities, 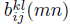, with cytonuclear inheritance. The A locus is biparentally inherited with µ the mutation probability from *A*_1_ to *A*_2_. The B locus is uniparentally inherited with _ the mutation probability from *B*_1_ to *B*_2_.

The top panel of Figure 3 shows the crossing time as a function of the mutation probability in the **B** locus, ν. Increasing ν increases the rate at which single and double mutants are created, aiding valley crossing. The bottom panel of Figure 3 shows the crossing time as a function of the ratio of the mutation probabilities at the two loci, *c*_*µ*_, while holding the average mutation probability, (µ + ν)/2 = µ(1 + *c*_µ_)/2, constant. When holding the average mutation probability constant the time to fixation is minimized when ν = µ because single mutant types are then equally frequent, increasing the chances they mate with one another to produce a double mutant by recombination. As *c*_*µ*_ departs from one, the mutation rate at one of the loci becomes small, causing those single mutants to become rare. The highly stochastic nature of the rare single mutant frequencies causes our semi-deterministic (Equation A3) and stochastic (Equation C5) approximations to underestimate the crossing time and, instead, the single mutants first reach mutationselection balance (Equation 8; dashed gray curve).

**Figure 3:**
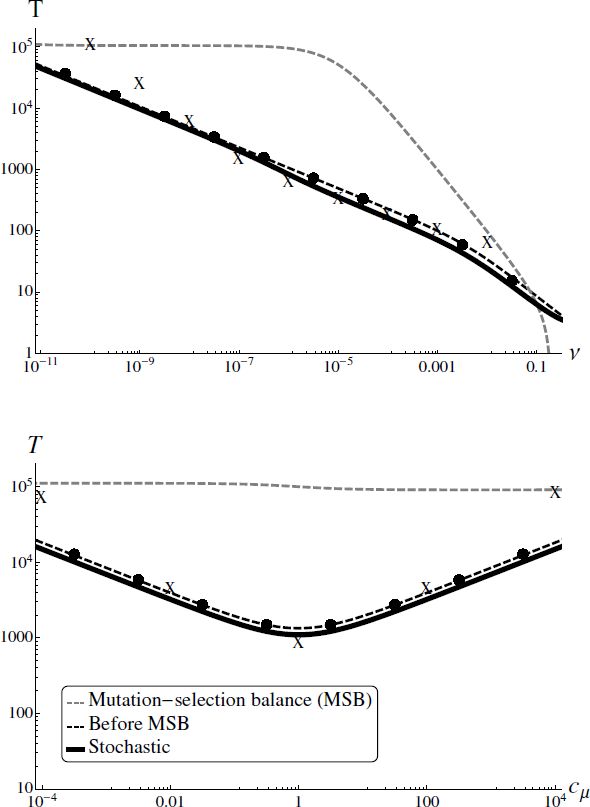
Expected number of generations until a double mutant begins to fix, *T*, as a function of **(top)** the mutation probability in locus **B**, ν, and **(bottom)** the relative mutability of the two loci, *c*_*µ*_ = *v*/µ. The top panel holds the mutation probability in locus A (µ = 5 x 10^−7^) constant while the bottom panel holds the average mutation probability [(µ + *v*)/2 = µ(1 + *c*_*µ*_)/2 = 5 x 10 ^7^] constant. The solid curves show the stochastic crossing time by recombination with neutral single mutants (Equation C5). The dashed curves show the semi-deterministic results when crossing occurs (*black*) before (Equation A3) and (*gray*) after (Equation 8) mutation-selection balance is first reached. The dots show the full semi-deterministic solution (numerical solution to Equation A1, including higher order terms, allowing both mutation and recombination to generate double mutants). The *X*’s are mean simulation results (Appendix D). Parameters as in Figure 1 **(bottom)**, except *w*_22_ = 2.01, which ensures *s*_22_ > 0 ∀ *c*_*µ*_.

Given that crossing occurs by recombination from standing variation (Equation 17), asymmetric mutation rates have little effect given a particular starting population (*i*_0_, *j*_0_). However, standing variation will also tend to vary in proportion to mutation rates, implying that uniparental inheritance will cause differences in the initial numbers of the two single mutants, which can have a large effect. Let *c*_*µ*_ now also determine the ratio of the initial numbers of single mutants, *c*_*μ*_ = *c*_*n*_ = *j*_0_/*i*_0_. Figure 4 shows the probability of crossing from standing variation as a function of *c*_*μ*_. When we hold *i*_0_ and s constant and increase *j*_0_ and ν (grey curve), the probability of crossing increases with *c*_*µ*_ as there are then more single mutants segregating. When we instead hold the total initial number of single mutants (*i*_0_ + *j*_0_) and the average mutation probability [(*μ* + ν)/2] constant (black curve), the probability of crossing is maximized at *c*_*μ*_ = 1 because the single mutants are then equally frequent and hence more likely to mate with one another and produce a double mutant through recombination.

**Figure 4:**
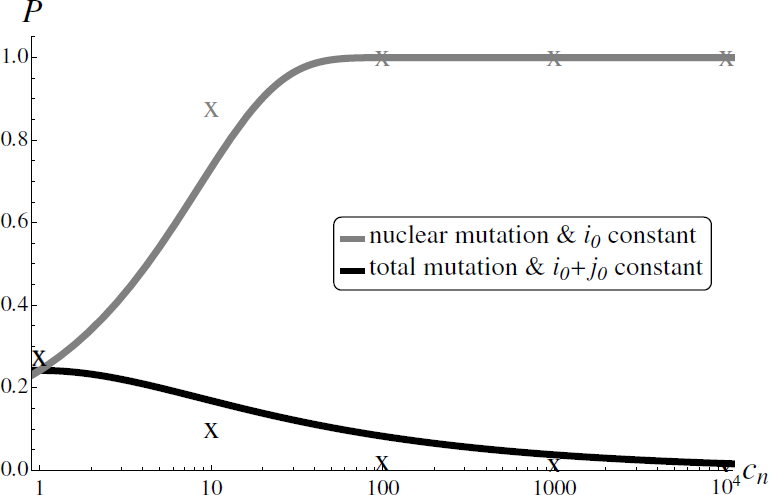
The probability, *P* = 1 − *u*, of crossing the valley given an initial stock of single mutants (with no further mutations from resident-resident matings) as a function of the ratio of the initial numbers of single mutants and mutation probabilities, *c*_*µ*_ = ν/_µ_ = *c*_*n*_ = *j*_0_/*i*_0_ (Equation 17). The *gray* curve holds the initial number of **A*_2_*B*_1_* (*i*_0_ = 100) and the mutation probability in the A locus (µ = 5 x 10^−7^) constant and varies the initial number of *A*_1_*B*_2_ (j_0_) and the mutation probability in the B locus (*v*). The *black* curve holds the initial number of single mutants (*n*_0_ = *i*_0_ + *j*_0_ = 200) and average mutation probability [(µ + ν)/2 = 5 x 10^−7^] constant. The X’s are simulation results (Appendix D). Other parameters as in Figure 3.

*2.3.3 Cultural inheritance*

Finally, we remove Darwinian selection, such that transmission bias alone determines the dynamics, and interpret the model in a cultural context. For the sake of exposition we consider only one simplified case of cultural transmission. Let trait combinations with only one new trait (*A*_2_*B*_1_ and *A*_1_*B*_2_) be inherited relative to the previous combination (*A*_1_*B*_1_) with probability q. Let the new combination of cultural traits (*A*_2_*B*_2_) be inherited relative to the previous combination with probability p. We are most interested in the case of a “transmission valley”, where the previous combination of traits is transmitted more effectively than mixed combinations of new and old (*q* < 1/2), but the all-new combination spreads even more effectively than the previous combination (*p* > 1/2). We assume that parental trait combinations can be broken up with probability *r* and mutation occurs with probability’s. The transmission probabilities are shown in Table 4.

**Table 4.**
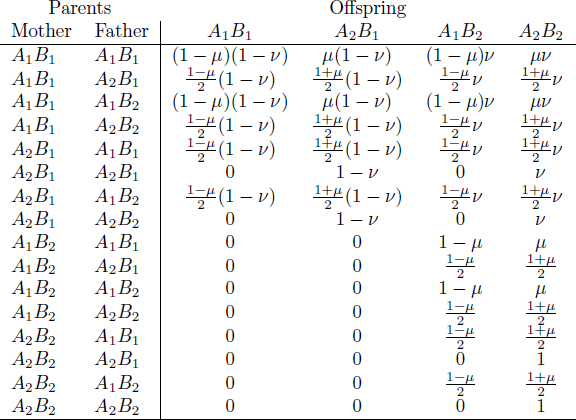
Transmission probabilities, 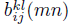, with cultural inheritance. Parental trait combinations are broken up with probability r, followed by biased transmission (*A*_2_*B*_1_ and *A*_1_*B*_2_ are passed down over *A*_1_*B*_1_ with probability q, *A*_2_*B*_2_ is passed down over *A*_1_*B*_1_ with probability *p*), and mutation with probability µ.

Figure 5 shows that the crossing time is substantially faster when the new combination of traits has a stronger transmission advantage (*T* decreases with p; compare thick curve with thin). Nevertheless, even combinations that are transmitted very effectively (thick curve) spread very slowly when their component traits are passed on poorly in the previous cultural background (*q* ≪ 1/2). In particular, the crossing time increases most quickly as *q* decreases from 1/2, demonstrating that slight biases in the transmission of the new traits when arising within the previous cultural background have a strong influence on the spread of new combinations of cultural traits, effectively preventing establishment if *q* ≪ 1/2.

**Figure 5:**
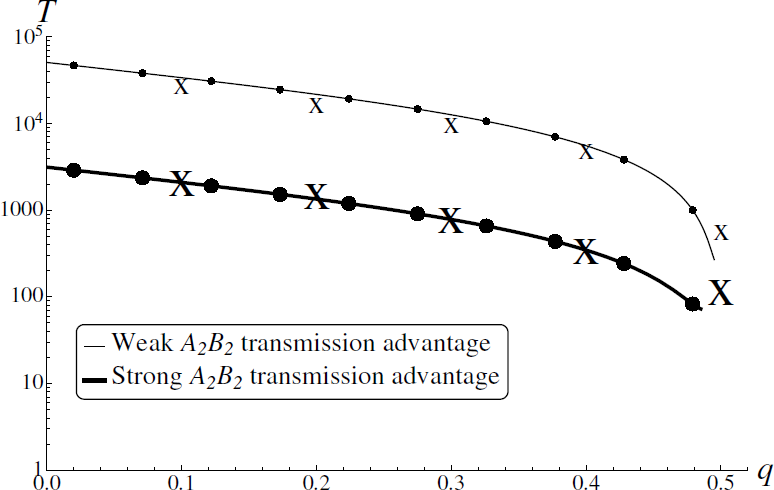
Expected number of generations until the new combination of cultural traits (*A*_2_*B*_2_) begins to fix, *T*, as a function of the probability of inheritance, *q*, of the new traits singly (*A*_2_ *B*_1_, *A*_1_*B*_2_) over the previous combination (*A*_1_*B*_1_). The curves show the estimate given mutation-selection balance is first reached (which assumes *A*_2_*B*_1_ and *A*_1_*B*_2_ are disfavoured, *q* < 0.5; Equation 8). The dots show the full semi-deterministic solution (numerical solution to Equation A1, including higher order terms, allowing both recombination and mutation to generate double mutants). The *X*’s are mean simulation results (Appendix D). The transmission advantage for the new combination of cultural traits is either weak (thin curves, small dots: *p* = 0.51) or strong (thick curves, large dots: *p* = 0.6). Parameters: N = 103, → = 10^−3^, *r* = 0.01, and *w*_21_ = *w*_12_ = *w*_22_ = 1.

Figure 6 shows the probability of crossing from standing variation. In this case, with such a large mutation rate, crossing can be more likely by mutation (Equation 15) than by recombination (Equation 17). Recombination has the added effect of breaking up the new combination of traits, reducing the probability of crossing. With a lower mutation rate crossing is most likely with moderate amounts of recombination (e.g., Figure 1). Figure 6 again shows that the transmission advantage of the new combination of traits (p; compare thick lines to thin) and slight biases in the transmission of new traits in the previous cultural background (q ≈ 1/2) greatly influence the probability that a new combination of cultural traits successfully spreads.

**Figure 6:**
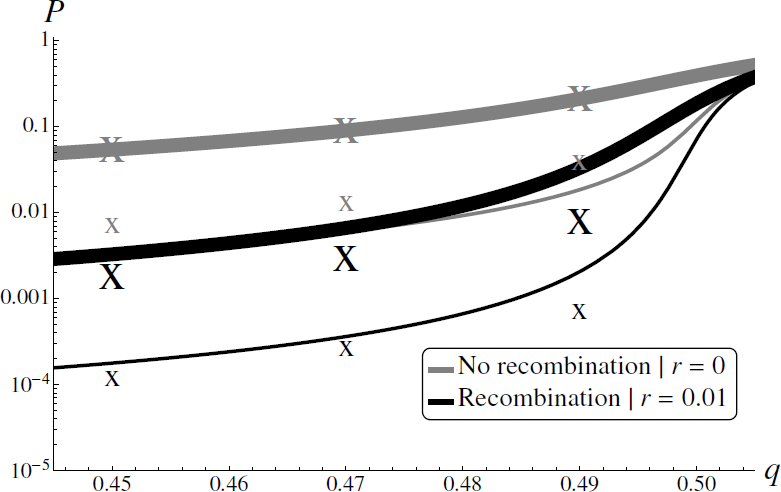
The probability, *P* = 1 − *u*, that the new combination of cultural traits (*A*_2_*B*_2_) fixes given an initial number of *A*_2_ *B*_1_ and *A*_1_*B*_2_ (with no further mutations from resident-resident matings) as a function of the probability of inheritance, *q*, of the new traits singly (*A*_2_*B*_1_, *A*_1_*B*_2_) over the previous combination (*A*_1_*B*_1_). The *grey* curves show the probability of crossing in the absence of recombination (r = 0; Equation 15), with a strong (thick curves: *p* = 0.6) or weak (thin curves: *p* = 0.51) transmission advantage for the new combination of cultural traits. The *black* curves show the probability of crossing by recombination only (r = 0.01; Equation 17). The *X*’s are simulation results (Appendix D). With such a large mutation probability, crossing can be more likely without recombination, which has the added effect of breaking apart the new combination. Parameters as in Figure 5 with *i*_0_ + *j*_0_ = 20 and *c* =1.

**3. Discussion**

Our results support the general consensus that, given reasonable population sizes and per locus per generation mutation rates, crossing a particular fitness valley by genetic drift is typically a slow and unlikely event (Crow and Kimura, 1965; Bengtsson and Bodmer, 1976; Lande, 1979; Hedrick, 1981; Walsh, 1982; Lande, 1985b; Michalakis and Slatkin, 1996; Phillips, 1996; Coyne et al., 1997). For example, with a per locus per generation mutation probability of *μ* = 10^−8^, a double mutant viability of *w*_22_ = 1.01, recombination between the two loci with probability *r* = 0.01, and a population size of *N* = 10^4^, the waiting time for a successful double mutant, in the best case scenario where single mutants are selectively neutral, is on the order of 10^7^ generations (Equation C5). As this is the typical age for living animal genera (Van Valen, 1973; Lande, 1979), we should not expect to see this fitness valley forded. Of course, with many potential fitness valleys across the genome, the chance that one of them is forded can become substantial.

By broadening previous treatments to allow for non-Mendelian inheritance, we have shown that a small amount of segregation distortion can greatly impact the chances of fitness-valley crossing. Of course, segregation distortion has a large impact because it provides a second level of selection (Sandler and Novitski, 1957), often acting like gametic selection (but see Hartl, 1970, 1977). When the *A*_2_ and *B*_2_ alleles are more likely to be passed down than the *A*_1_ and *B*_1_ alleles, respectively, in matings between single mutants and residents, the depth of the valley is effectively reduced and hence crossing is much more likely. For example, when single mutants have a relative viability of *w*_*m*_ = 0.95, the mutation rate is *µ* = 10^−8^, double mutants are weakly favoured (*w*_22_ = 1.01), and we begin with one single mutant (n_0_ = 1) in a population of size N = 10^4^, in the absence of recombination 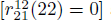 and segregation distortion (*k*_*ij*_ = 0), the probability of crossing is on the order of 10^−9^ (Equation 15). With a 5% distortion in favour of *A*_2_ and *B*_2_ alleles (*k*_21_ = *k*_12_ = 0.05) the single mutants are effectively neutral and the probability increases seven orders of magnitude to 10^−2^. And with a 10% distortion the single mutants are selectively favoured and the double mutant fixes with probability 0.25.

Segregation distortion, in the form of meiotic drive, has often been implicated as a force that could help fix underdominant chromosomal rearrangements (Sandler and Novitski, 1957; Bengtsson and Bodmer, 1976; Hedrick, 1981; Walsh, 1982; Faria and Navarro, 2010). Chromosomal rearrangements, such as translocations and inversions, are often fixed in alternate forms in closely related species (White, 1978; Coyne, 1989; Faria and Navarro, 2010). Because heterokaryotypes typically have severely reduced fertility (Sandler and Novitski, 1957; Lande, 1979), such rearrangements are thought to promote rapid speciation (stasipatric speciation; White, 1978, but see Faria and Navarro, 2010; Kirkpatrick, 2010). The trouble is explaining how such rearrangements originally increase in frequency when they are so strongly selected against when rare (Navarro and Barton, 2003; Kirkpatrick, 2010). Meiotic drive provides one possible answer. Our results can be used to investigate valley crossing with chromosomal rearrangements by assuming A and B are homologous chromosomes, with *A*_2_ and *B*_2_ being the novel chromosomes, and *A*_2_ *B*_1_ and *A*_1_ *B*_2_ interchangeable. For example, with free recombination 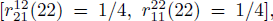, a 5% viability reduction in heterokaryotypes (*w*_*m*_ = 0.95), no meiotic drive 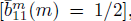, a very beneficial mutant homokaryotype (*w*_22_ = 2.5), and a spontaneous chromosome mutation rate of *µ* = 10^−3^ (Lande, 1979), when starting with one copy of each mutant chromosome (*i*_0_ = 1, *c*_*n*_ = 1) in a population of size *N* = 10^4^, Equation (17) gives a 0.4% chance of fixing the mutant homokaryotype. When the mutated chromosome has a 70% chance of being passed down in matings with residents, a relatively weak amount of drive (Sandler and Novitski, 1957), the chance of crossing increases two orders of magnitude, to nearly 75%.

Here we have shown that, for a given number of single mutants, the chance of crossing a valley by recombination is best when the two single mutant types are at equal frequencies. This is an important factor when the mutation rates in A and B are highly asymmetric. One instance where this asymmetry is likely is when one locus (say B) is in the mitochondrial genome, and is passed down maternally, while the other (say A) is in the nuclear genome, and is passed down biparentally. Mutation rates in the mitochondria can be orders of magnitude higher than in the nucleus (Linnane et al., 1989). With 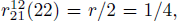, *N* = 10^4^, *w*_22_ = 2, 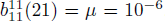, and neutral single mutants, when the mutation rates in A and B are equal 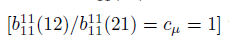 the crossing time is 40, 000 generations. When the mutation rate in **B** is two orders of magnitude larger (*c*_*µ*_ = 100) the waiting time is reduced to 2, 500 generations. But when the average mutation rate 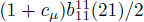 is held constant, the asymmetrical mutation rates instead hinder crossing, increasing the crossing time to nearly 120, 000 generations.

By expanding a mathematical model of fitness-valley crossing beyond symmetrical Mendelian inheritance we gain insight into transitions between alternate stable states in non-genetic systems, such as culture. As mentioned in the introduction, culture may often exhibit alternate stable states; here valley crossing corresponds to a shift between alternate combinations of cultural ideas or practices (e.g., the demographic transition; Borgerhoff Mulder, 1998). The valley is a “transmission valley”, created by new cultural traits that are transmitted effectively in concert but poorly when arising individually within the previous cultural background. In this case our simplified example above demonstrates that, given that the component pieces are not passed on too poorly in the previous cultural background, the probability that a new set of practices or ideas becomes pervasive in society is greatly improved by its transmission advantage over the previous set. Valley crossing might also be relevant in the context of gene-culture coevolution, where one trait is cultural and the other genetic. For instance, the ability to absorb lactose as an adult is largely genetically determined and is positively correlated with the cultural practice of dairy farming, reaching frequencies over 90% in cultures with dairy farming but typically remaining less than 20% in cultures without (Feldman and Laland, 1996). If, as seems reasonable, the ability to absorb lactose as an adult has a cost in the absence of dairy farming and the cultural practice of dairy farming has a cost when adults are unable to absorb lactose, then the transition from non-pastoralist non-absorbers to pastoralist absorbers may represent another example of fitness-valley crossing outside the purely genetic arena. We have used our generalized model to begin to explore cultural transitions, but it should be emphasized that we neglect oblique and horizontal transmission, common features of cultural evolution (Cavalli-Sforza and Feldman, 1981) and likely components of the demographic transition (Ihara and Feldman, 2004). Generalizing models of fitness-valley crossing further to include oblique and horizontal transmission would improve insight into cultural transitions.

We have incorporated transmission bias in a model of multi-locus fitness-valley crossing. This allows us to investigate fitness-valley crossing in new scenarios, such as in genetic systems with segregation distortion and/or uniparental inheritance. Segregation distortion acts as a second level of selection and therefore can greatly help or hinder fitness-valley crossing, especially when crossing is otherwise unlikely. Uniparental inheritance will often imply asymmetric mutation rates, which in turn lead to unequal frequencies of single mutants, and therefore, all else being equal, a lower probability of fitness-valley crossing by recombination. However, uniparental-inherited cytoplasmic elements tend to have increased mutation rates, which helps crossing. Generalizing transmission also allows us to begin to extend the theory of valley crossing to nongenetic systems, such as culture. Despite component traits being passed on poorly in the previous cultural background, we find that small advantages in the transmission of the new set of cultural traits will greatly facilitate a cultural transition. While crossing a deep fitness valley is difficult under Mendelian inheritance, it can be easier when Mendel is left behind.

## Acknowledgements

### 4. Acknowledgements

We thank Eva Kisdi, Stefan Geritz, Helene Weigang, and the labs of S.P. Otto, Michael Whitlock, Michael Doebeli, and Christoph Hauert for helpful discussions. We thank Mark Kirkpatrick for suggesting the simplicity of considering crossing times from mutation-selection balance. Funding provided by Natural Sciences and Engineering Research Council (Canada) Discovery (SPO) and CGS-D (MMO) grants.

## A. Dynamic single mutants

Here we calculate the waiting time for a successful double mutant to arise, starting from a population that is composed entirely of residents. While single mutants are far from mutation-selection balance, |*s*_*ij*_*t*| < 1 ∀ *i* ≠ *j*, we can write *x*_22_ as a function of *t* by replacing *x*_*ij*_ in Equation (4) with the appropriate *x*_*ij*_ in Equation (3). With rare mutation, rare single mutants, and weak selection on single mutants, the expected number of generations until a successful double mutant appears, *T*, solves

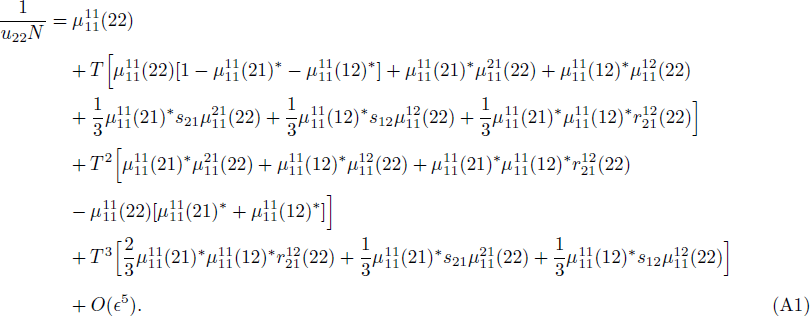

The O(ε^5^) terms disappear and the equation is exact when single mutants are neutral, *s*_*ij*_ = 0. Otherwise, with selection against single mutants, the higher order terms can only be ignored as long as the crossing time, *T*, is much smaller than the inverse of the selection coefficients, *s*_21_ and *s*_12_. Because Equation (A1) is a cubic in T, its solution is cumbersome (see supplementary *Mathematica* file). Here we examine two scenarios which give more interpretable approximations for T.

Without selection on single mutants (*s*_21_ = *s*_12_ = 0) and without recombination from single mutants to double mutants 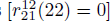 the *T*^3^ term in Equation (A1) vanishes. In addition, if the crossing time T is long, the dominant term is the one proportional to *T*^2^. Solving for *T* from this term alone gives

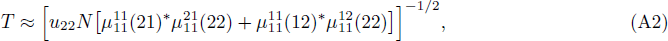

where we have ignored double mutants arising instantaneously 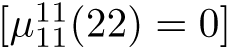. Equation (A2) shows that the crossing time without selection on or recombination among single mutants is roughly proportional to *N*^−1/2^ generations. The crossing time decreases with N because increasing N increases the per generation input of mutations. Holding mutation input 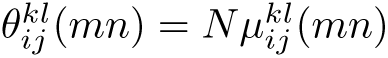 constant, the crossing time becomes proportional to *N*^1/2^. When the single-mutation transmission probabilities are equal 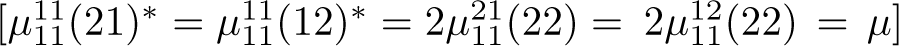 and we calculate the first appearance of any double mutant (successful or not; *u*_22_ = 1), the expected time until the first double mutant appears simplifies to the neutral genetic case without recombination, 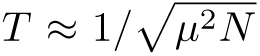 (equation 8 in Christiansen et al., 1998). Equation (A2) clarifies the role of the various, *potentially different*, mutation probabilities 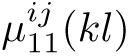 on the time until the first double mutant, while also allowing us to ignore double mutants that are lost.

When there is recombination between single mutants to produce double mutants 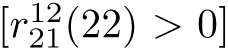 and the crossing time, *T*, is smaller than the inverse of the selection coefficients, *s*_12_ and *s*_21_, the dominant term in Equation (A1) is proportional to *T*^3^. This term is positive when recombination is frequent relative to selection against single mutants. Again, if the time *T* is long we can use this term alone to approximate *T*, which gives

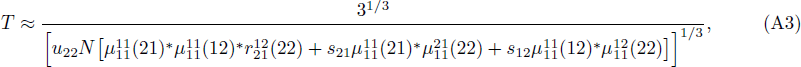

where we have once again ignored the instantaneous production of double mutants. Notice that, for a given mutation input 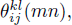 when there is recombination between single mutants, the crossing time is roughly proportional to *N*^1/3^ generations (rather than *N*^1/2^ generations without recombination), implying that recombination between single mutants tends to shorten the expected time until the first (successful or unsuccessful) double mutant arises. However, because recombination can also occur between residents and double mutants (reducing *u*_22_) Equation (A3) shows that the waiting time until the first *successful* double mutant is minimized at intermediate levels of recombination.

Equation (A3) reduces to 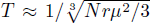 (equation 9 in Christiansen et al., 1998) when we ignore the weak selection against single mutants (*s*_*ij*_ = 0), there is equal mutation probability at each locus 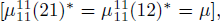, and we wait until the first double mutant appears, successful or not (*u*_22_ = 1). Once again our analysis clarifies the role of the various, potentially different, mutation probabilities 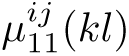 on the waiting time until the first successful double mutant. Equation (A3) also allows (weak) selection on single mutants and incorporates transmission bias, which we explore more fully in the main text.

Figure *A*_1_ compares the approximations derived here (Equations *A*_2_ and *A*_3_) with that derived in the text assuming mutation-selection balance is reached before crossing (Equation 8). The approximations given by Equations (A2) and (A3) break down as the depth of the valley (**δ** = −*s*_21_ = *s*_12_) increases such that the crossing time becomes long, T > 1/**δ**.

**Figure A1:**
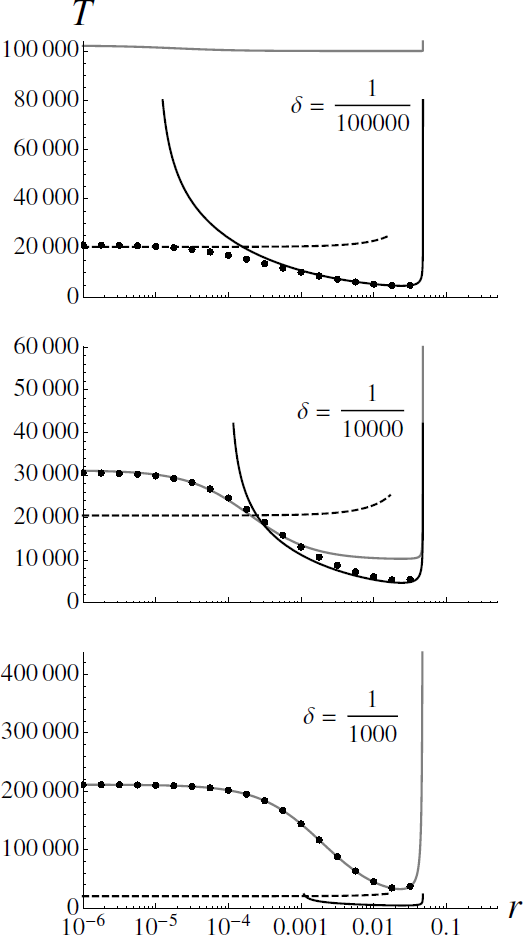
Expected number of generations until a double mutant begins to fix, *T*, as a function of recombination, *r*, given (*gray*) mutation-selection balance is first reached (Equation 8) or mutation-selection balance is not reached and *(black, solid*) crossing can occur by recombination (Equation A3) or *(black, dashed*) crossing occurs by mutation only and − *S*_21_ = − *S*_12_ = 6 = 0 (Equation A2). The dots show the full semi-deterministic solution (numerical solution to Equation A1, including higher order terms, allowing both recombination and mutation to generate double mutants). The mutation-selection balance estimate (*gray*) performs better than the dynamic estimates (*black*) when *δ T* > 1, and vice-versa. Parameters: symmetrical, Mendelian inheritance with N = 10^5^, *s*_22_ = 0.05, and → = 5 x 10^−7^ (see supplementary *Mathematica* file).

## B. Diffusion approximation

Here we take the large population limit (N → ∞), scale time such that one unit of time in the scaled diffusion process 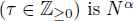 generations in the unscaled Markov process (Δ*t* = *τN*^α^) and define new frequency parameters 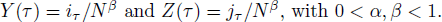

We are concerned with three quantities for each variable ΔY and ΔZ. The first is the infinitesimal mean

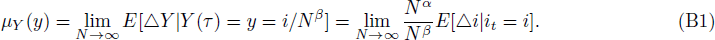

The second quantity is the infinitesimal variance

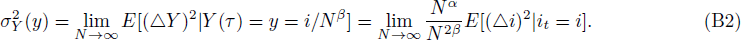

And the third quantity of interest is a higher (*n* > 2) infinitesimal moment

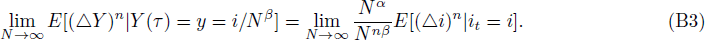

We can similarly calculate 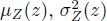, and a higher moment in ΔZ.

The final quantity of interest is the scaled “killing rate”

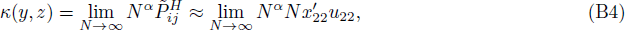

where the approximation assumes 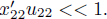

For the Markov chain to converge to a diffusion process as *N* → ∞ to we require: 1) μ_*Y*_ (*y*) and μ_*Z*_ (*z*) to be finite; 2) 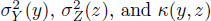 to be positive and finite; and 3) some higher moment (in both Δ*Y* and Δ*Z*) to be equal to zero (Karlin and Taylor, 1981). We first take a hint from the genetic case (Christiansen et al., 1998) and scale transmission probabilities as

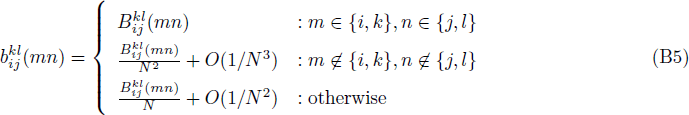

In the genetic case this can be interpreted as making the probability of mutation proportional to the inverse of population size *µ* = *B*/*N*. Then, as *N* → ∞ mutation probability decreases (*µ* → 0), such that mutation input *B* = *Nμ* is constant. This prevents the process from taking large jumps in frequency space, which violate the diffusion process (Karlin and Taylor, 1981).

In order for the transmission parameters to satisfy the logical constraint 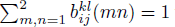 the diffusion also requires, as *N* → ∞, that

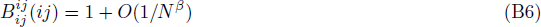

and

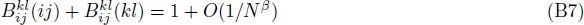

when either {*i* ≠ *k*, *j* ≠*l*} or {*i* = *k*, *j* ≠ *l*}. In words, the sum total mutation probability for parents *A*_*i*_*B*_*j*_ and Ak Bi; must be relatively small, at most on the order of 1/*N^β^*.

Finally, our approximation requires weak selection, relative to *w*_11_ = 1. In particular, total selection on single mutants must be weak, on the order of 1/*N^β^*,

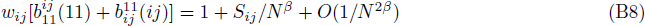

for i ≠ j, where *s*_*ij*_ is the scaled selection strength. And selection on double mutants must also be weak, such that

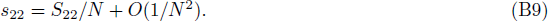

With the above assumptions (Equations B5-B9) the Markov chain converges to a diffusion process as N **→ ∞** when

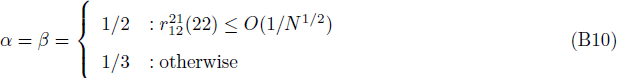

This scaling implies that if recombination between single mutants to make double mutants 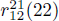 is less likely that mutation (which is on the order of *N*^−1/2^; Equation B6), then the time until the process is killed scales with *N*^1/2^. Meanwhile, if recombination is more likely than mutation the killing time scales with *N*^1/3^. These results align with our semi-deterministic analysis (Equations *A*_2_ and *A*_3_).

When **α** =**β** the infinitesimal variances are 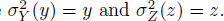 The infinitesimal means and the killing term depend on the probability of recombination. When recombination is rare the single mutants are expected to reach higher frequencies and therefore have a greater influence on the dynamics. To simplify, when recombination is rare 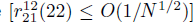 we assume weak transmission bias for residents mating with single mutants 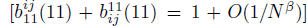 and for single mutants mating with each other 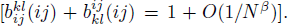 We further assume weak viability selection on single mutants, *w*_*ij*_ = 1 *O*(1/*N^β^*), regardless of recombination. The infinitesimal mean is then always

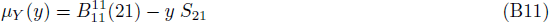

and similarly for μ_*Z*_(*z*). The first term, 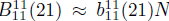, describes mutation to single mutants in resident-resident matings and the second term, with *S*_21_ ≈ *s*_21_N^β^, describes the removal of single mutants by selection (both through transmission bias when mating with the resident and survival).

The killing terms are

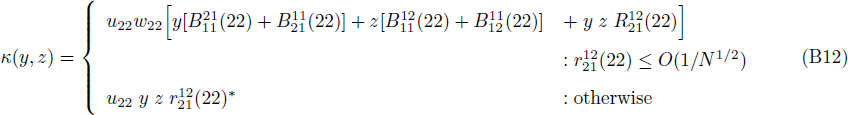

where 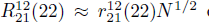 describes a (low) probability of recombination. The first line shows that the process can be killed by mutations in single mutants that mate with residents 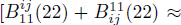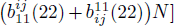 or by rare recombination between single mutants to produce double mutants 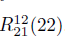. When recombination is more likely than *N*^−1/2^ the process is essentially always killed by recombination 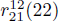 (second line in Equation B12).

## C. Stochastic crossing times

*Neutral single mutants without recombination.* With no chance of recombination from single mutants to double mutants 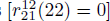 we have scaling parameter **β**= 1/2. Then, without selection on single mutants (*s*_21_ = *s*_12_ = 0) and with some mutational symmetry between the two loci 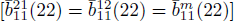, the single mutants are equivalent and we can concern ourselves with only their sum, ξ = *y* + *z*. Letting m be either single mutant type (*m* = 21 or 12), Equation (12) reduces to

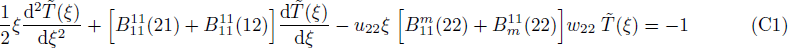

where 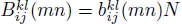.

When there are an infinite number of single mutants a successful double mutant is produced immediately, giving one boundary condition 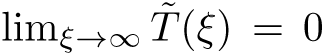 The second boundary condition is 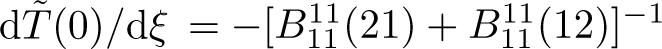, which can be derived directly from Equation (C1) by setting ξ = 0 (see appendix A in Christiansen et al., 1998, for a more complete derivation).

The solution to the boundary value problem, evaluated at ξ = 0, corresponding to the expected number of generations until a successful double mutant arises when beginning with only residents, 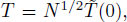, is then

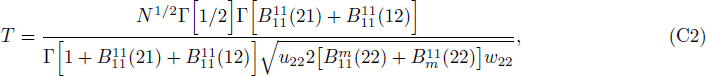

where Г[·] is the gamma function. Setting mutation probabilities equal 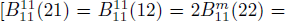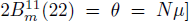 reduces Equation (C2) to the neutral genetic case (equation 27 in Christiansen et al., 1998) divided by 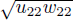 because we census after selection and consider double mutant fixation. By separating the various mutational terms our analysis clarifies that, while the crossing time is inversely proportional to the mutation probability from residents to single mutants, it is inversely proportional to the *square root* of mutation probabilities from single mutants to double mutants. The crossing time is therefore increased much more by a reduction in mutations from residents to single mutants than it is by a reduction in mutations from single mutants to double mutants.

When mutations from residents to single mutants 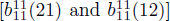 are rare, an approximation for the crossing time, in terms of our unscaled parameters, is

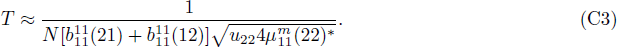

Increasing the mutational supply of single mutants 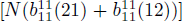 or the probability of mutation from single mutants to successful double mutants 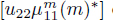 decreases the amount of time we expect to wait before a successful double mutant arises. Holding mutation input, 0, constant, Equation (C3) shows that the crossing time without recombination is roughly proportional to *N*^1/2^ generations, aligning with the semi-deterministic analysis (Equation A2) and indicating that, for a given mutational input, genetic drift increases the speed at which fitness valleys are crossed.

*Neutral single mutants with recombination.* With recombination the scaling parameter is **β**= 1/3. We can reduce and solve Equation (12) with recombination when the frequencies of single mutants remain proportional to one another, such that we need follow only *c_μ_ y* = *z* = ξ, where *c*_*µ*_ is a constant. This requires mutation input 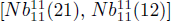 to be large enough to make the dynamics of y and z relatively deterministic. We further assume no selection on single mutants (*s*_21_ = *s*_12_ = 0). We then have *c*_*μ*_ y = z for all time, *t*, when the ratio of mutation probabilities is 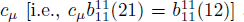 and we begin with *c*_µ_*y*(0) = *z*(0). Equation (12) then collapses to

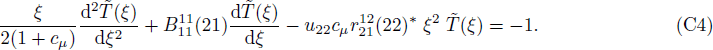

The boundary conditions are 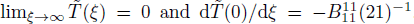. The solution to the boundary-value problem, evaluated at ξ = 0, in units of generations, 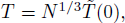, is

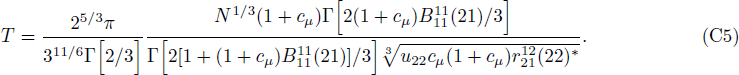

Letting 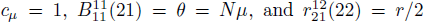 reduces Equation (C5) to the neutral genetic case (equation 30 in Christiansen et al., 1998) divided by 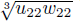 because we census after selection and consider double mutant fixation. Our result extends the insight of Christiansen et al. (1998) by allowing the frequencies of single mutants to differ, *c*_*µ*_ ≠ 1. Holding average mutation input 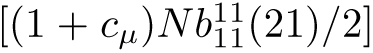 constant, Equation (C5) shows that the crossing time is minimized when there are equal numbers of the two single mutants (*c*_µ_ = 1) and increases as the asymmetry grows. This occurs because recombination is most effective in creating double mutants when the single mutants are equally frequent.

Converting the full solution back in terms of our unscaled parameters and letting the mutation probability 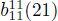 be small, we have the approximation

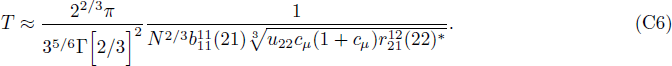

Holding mutation input 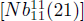 constant, Equation (C6) shows that the crossing time is roughly proportional to *N*^1/3^ generations, aligning with the semi-deterministic analysis (Equation A3).

## D. Stochastic simulations

We performed stochastic simulations to verify our analytical and numerical results. Simulation code is supplied in the supplementary *Mathematica* file. Briefly, we performed random multinomial sampling of genotypes with frequency parameters given by Equation (1) and transition probabilities defined in Tables 2-4. Crossing time simulations ended on double mutant fixation and the generation in which this occurred was recorded as the crossing time. Crossing time was averaged over all trials (103 trials in Figure 1, 102 trials in Figure 3 and 5). Crossing probability simulations ended on resident or double mutant fixation and the genotype which fixed in each trial was recorded. The crossing probability was calculated as the fraction of trials in which the double mutant fixed (10^3^ trials in Figure 2 and 4, 10^5^ trials in Figure 6).

